# The physical origin of ageing at biomolecular condensate interfaces

**DOI:** 10.64898/2026.03.05.709844

**Authors:** Alejandro Castro, Juan Luengo-Márquez, Andres R. Tejedor, Marcell Papp, Paolo Arosio, Alberto Ocaña, Rosana Collepardo-Guevara, Ignacio Sanchez-Burgos, Jorge R. Espinosa

**Author notes:** To whom correspondence should be sent.

## Abstract

The interface separating a biomolecular condensate from its surroundings is increasingly recognised as a unique physical and chemical environment that actively regulates condensate behaviour. It controls surface tension, molecular organisation and chemical reactivity, and is also where pathological ageing often begins, with several proteins nucleating cross-β-sheet fibrils preferentially at condensate surfaces. But does ageing emerge there because of protein-specific chemistry, or a more general physical mechanism? Using coarse-grained molecular dynamics simulations, we show that cross-β-sheet nucleation generically occurs preferentially at condensate interfaces across systems of increasing chemical complexity, from simple homopolymers to patterned amphiphilic sequences. In all systems studied, interface-driven ageing has a common physical origin and can be further modulated, but not driven, by protein sequence. Specifically, terminal domains at the condensate interface gain mobility while remaining densely packed and mutually aligned, creating favourable conditions for β-sheet nucleation. Ageing is amplified in amphiphilic sequences, where hydrophilic regions act as surfactants facing the dilute phase, while aggregation-prone hydrophobic domains cluster just beneath the surface, concentrating the nucleation there. This surfactant-like organisation lowers the interfacial free energy, which falls systematically as sequence asymmetry grows. By linking molecular interactions to mesoscale behaviour, these results cast the interface as a central regulator of ageing and reveal a sequence-independent physical mechanism for the liquid-to-solid transition underlying pathological aggregation in biomolecular condensates.

## I. INTRODUCTION

Cells organise their biochemistry not only through membrane-bound organelles^1^ but also through membraneless compartments known as biomolecular condensates^2–6^. Ubiquitous across the cell—from stress granules to the nucleolus and Cajal bodies^7–11^—condensates concentrate proteins and nucleic acids to control where and when biochemical reactions occur. Because they can assemble and disassemble dynamically, they provide a tunable mean of spatiotemporal regulation^12–14^.

Condensates are postulated to form by phase separation of proteins, nucleic acids and other macromolecules engaged in multivalent, transient interactions^5,15^. Such interactions are frequently mediated by RNA through specific and non-specific protein–RNA contacts^8,16–18^ and complemented by protein low-complexity domains (LCDs) that engage in weak, multivalent protein–protein interactions^4,19,20^. Condensate properties are finely regulated by multiple external factors, including, temperature, salt and pH^21–25^. They are also vulnerable to undergoing liquid-to-solid transitions linked to neurodegeneration^26,27^. These transitions underlie disorders such as ALS, where mutations alter the phase behaviour of RNA/DNA-binding proteins including TDP-43 and FUS^28–31^; Parkinson’s disease, linked to α-synuclein fibrillation^32^; and dementias driven by Tau misfolding^33–35^.

The conversion of a functional, liquid-like condensate into a pathological solid-like state—termed ageing—often proceeds through inter-protein structural transitions that generate inter-protein cross-β-sheet motifs^36^. Their progressive accumulation strengthens the intermolecular network of the condensate, shifting condensates from transient, liquid-like behaviour to kinetically trapped solids^37,38^ and steadily slowing molecular diffusion as β-sheet content rises^39–41^. Increasingly, this ageing is traced to the condensate interface^42^. Interfacial properties—surface tension^43,44^, zeta potential^45^ and molecular adsorption^46^—govern condensate size^47,48^, stability, and interactions with cellular structures such as autophagosomes^49^, microtubules^50^ and endocytic pits^51^, while also shaping molecular orientation and reactivity at the interface^52^. Consistent with this emerging view, the liquid-to-solid transition in FUS condensates has been reported by us to initiate at the droplet boundary^53,54^. Similarly, in hnRNPA1 condensates, cross-β-sheet formation during ageing occurs preferentially at the interface^40^, as well as α-synuclein forms fibrils which accumulate at condensate surfaces^32,55^.

Yet why ageing preferentially begins at condensate inter-faces remains unresolved. Proteins such as FUS, hnRNPA1 and α-synuclein each possess distinct aggregation-prone sequences, making it difficult to determine whether interfacial ageing arises from protein-specific chemistry or from a more general physical property of condensate surfaces. Experiments alone cannot readily disentangle these contributions because both are inherently coupled in real proteins. Simulations, by contrast, allow sequence chemistry to be systematically removed or reintroduced, providing a route to isolate the chemical and physical principles that govern ageing at condensate interfaces.

Here, we use coarse-grained molecular dynamics (MD) simulations, at both equilibrium and non-equilibrium, to map where inter-protein structural transitions occur within condensates and to identify the factors that drive them^39,54,56^. We deliberately begin from a minimal polymer-physics model of a protein LCD. By stripping away sequence-specific chemistry, this model isolates the physical contribution to ageing while recapitulating essential condensate behaviour, including fusion dynamics^41^, disorder-to-order conformational transitions^57^, and sequence-dependent stability^58–61^. Within each protein sequence, we define sites capable of structural transitions (Fig. 1a) that emulate low-complexity aromatic-rich kinked segments (LARKS), which are known to promote intermolecular β-sheet formation^62^. We then progressively introduce chemical heterogeneity, from purely hydrophobic to amphiphilic LCDs, and characterise condensates before and after ageing through local density fluctuations, conformational ensembles, intermolecular rearrangements, monomer diffusion, contact-frequency maps and surface tension. Across all sequences, structural transitions nucleate preferentially at the condensate interface. Crucially, we find that, in all cases, interface-driven ageing arises from a common physical mechanism: at the interface, molecules remain mobile while experiencing strong local density fluctuations and enhanced inter-protein alignment, creating favourable conditions for inter-protein βsheet formation. Sequence chemistry modulates the strength of this effect but does not determine where ageing begins. The distinctive chemistries of proteins such as FUS^53,54^, hnRNPA1^40^ and α-synuclein^32,55^ therefore amplify or suppress interfacial ageing rather than create it. Therefore, our results suggest that condensate ageing is governed by generic physical features of the interface, including surface tension, interfacial composition, and the local coexistence of mobility, density fluctuations and molecular alignment, rather than by sequence-specific chemistry alone. This identifies ageing at condensate interfaces as a general physical mechanism across diverse condensate-forming proteins implicated in neurodegeneration and points towards interfacetargeted strategies to control pathological aggregation.

**Figure 1:**
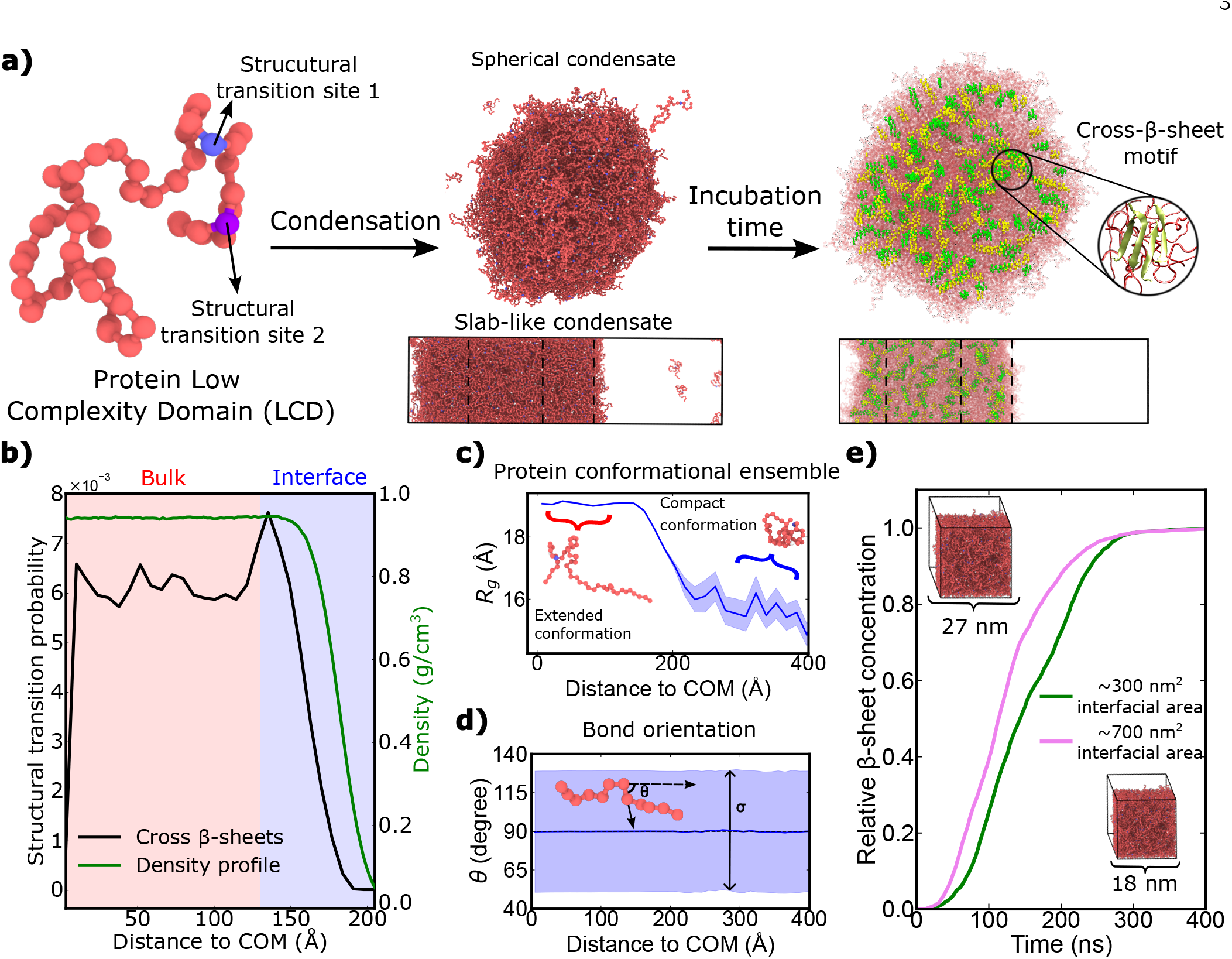
(a) Schematic representation of our minimal protein low-complexity domain (LCD) model which is able to undergo phase-separation and subsequent evolution, upon incubation time, into solid-like states driven by the accumulation of inter-protein β-sheets. The LCD sequence is formed by 50 homotypic residues, where beads emulating LARKS are highlighted in blue and purple. The renders depict the organisation of the LCD within spherical and slab-like condensates (centre) and the cross-β-sheet structures formed over time (right). (b) Cross-β-sheet structural transition probability (black) and density profile (green) as a function of distance from the condensate centre-of-mass (COM). (c) Radius of gyration (R*_g_*) as a function of distance from the condensate COM. Rendered images show representative conformations observed at different distances. The error is represented by a shaded region. (d) Average angle between the bond vector connecting directly linked residues and the direction normal to the interface (θ), shown as a function of distance from the condensate COM. The width of the θ distribution at each distance (σ) is indicated by the shaded region. (e) Time evolution of cross-β-sheet concentration for two condensates with different interfacial areas but constant number of LCDs. Rendered images illustrate the difference in box dimensions between both simulations to control the surface to volume ratio.

## II. RESULTS AND DISCUSSION

### A. A Minimal Low-Complexity Domain Model for Condensate Formation and Ageing

We employ an off-lattice implicit solvent protein LCD model based on the Mpipi Recharged sequence-dependent force field^63^, in which each residue is represented as a spherical bead in implicit solvent (see Section S1 of the Supplementary Material, SM). Intermolecular hydrophobic interactions are approximated using the Wang–Frenkel potential^64^, while chain connectivity is enforced through a bond harmonic potential. Our MD simulations are performed in LAMMPS^65^ in combination with an ageing algorithm developed by us^22,39,54,66–68^ that dynamically mimics intermolecular β-sheet transitions in coarse-grained protein models when multiple LARKS (i.e., structural transition sites depicted in blue; Fig. 1a) fall within a prescribed cut-off distance. Upon satisfying this distance and coordination criterion^54,67^, pairwise interactions between the involved residue types are rescaled from weak, transient LCD-like contacts to strong β-sheet-like interactions, informed by all-atom calculations^39,54,67^ (see Section S1 of the SM for details). To simulate phase-separation, we focus on two classes of condensate architectures: spherical droplets and slab-like geometries, as illustrated in Fig. 1a. Slab simulations enable the calculation of phase diagrams in the temperature–concentration plane^22^ (see Section SII of SM for further details), while spherical condensates, consisting of a stable droplet embedded within a cubic simulation box, permit the evaluation of high surface to volume ratios, as experimentally relevant for condensate nucleation events^69,70^.

We explore the impact of LCD intermolecular interactions in condensate ageing by designing three representative sequences recapitulating small differences in the net interaction strength. The first protein LCD consists of uniform interactions along the entire sequence, corresponding to a homopolymer. The second adopts a block-copolymer architecture with a hydrophobic region characterized by enhanced intermolecular interactions and a hydrophilic region with scale-down inter-protein interactions. Finally, the third sequence follows a similar block-copolymer arrangement but with pairwise interactions tuned to accentuate the contrast between the hydrophobic and hydrophilic segments. These amphiphilic LCDs therefore provide a more faithful approximation of real LCD phase behaviour forming biomolecular condensates and ultimately transitioning into solid-like kinetically trapped states^38^.

### B. Ageing emerges at condensate interfaces even for chemically uniform homopolymers

We begin with the simplest case—a chemically uniform homopolymer LCD—to ask whether structural transitions localise at the interface in the absence of any sequence patterning. We first compute the temperature– concentration phase diagram for the homopolymer LCD (hereafter referred to simply as LCD; Fig. SII) to determine the coexistence regime between condensed and dilute phases (Fig. 1a). From these simulations, using the Direct Coexistence method^71^, we estimate a critical solution temperature of T_c_ = 278 ± 2 K. The system is then equilibrated at T/T_c_ = 0.92, and we perform non-equilibrium simulations of condensate ageing using our dynamical algorithm^39,54^, which mimics inter-protein β-sheet structural transitions when at least 4 LARKS are found in a high-density local fluctuation, inducing an enhancement of the interaction strength equivalent to those found upon a LARKS disorder-to-order transition^62,67,72^. Within these simulations, inter-molecular interactions between residues capable of undergoing structural transitions (blue and purple beads in Fig. 1a) are dynamically switched over time, mimicking disorder-to-order structural transitions^62^. We characterize condensate ageing by spatially tracking through the MD trajectory the position of the spontaneous structural transitions relative to the condensate centre of mass (COM). The results, shown in Fig. 1b, reveal a pronounced maximum in the probability of structural transitions occurring near the condensate interface (black curve), averaged over ten independent simulation trajectories. The interfacial region of interest considered in this study is defined as extending two molecular diameters from the point at which the density profile reaches the bulk value (i.e. two times σ) on both sides. While the presented data in Fig. 1b corresponds to slab-like condensates with an interfacial area of 324 nm^2^, similar trends are observed for different condensate surface to volume ratios, as well as for spherical condensates, as reported in Sections SIII and SIV of the SM. For spherical condensates, however, the smaller volume of the inner shells results in limited sampling statistics when normalized by volume.

Remarkably, we find that our minimal LCD model recapitulates the enhanced probability of structural transitions at the condensate’s interface, thereby capturing a key feature experimentally reported for phase-separating RNA-binding proteins such as FUS or hnRNPA1^40,53^. While this interfacial initiation has been observed experimentally and in our own earlier sequence-specific simulations, its physical origin has remained unresolved. The minimal model simulations in this work allows us to uncover such origin. This is our first central result: interfacial ageing arises even in a homopolymer entirely devoid of chemical patterning, establishing that sequence-specific chemistry is not required to localise ageing at the interface.

Understanding the mechanistic origin of this interfacial preference is crucial for elucidating how ageing driven by cross-β-sheet transitions unfolds in biomolecular condensates. Given that our model does not incorporate chemical heterogeneities encoded across the protein sequence, we hypothesize that the observed maximum in transition probability may arise from the influence of the interface on the LCD conformational ensemble. To test this hypothesis, we analyze the LCD radius of gyration (R*_g_*) (Fig. 1c) and bond orientation as a function of the COM distance from the condensate’s core (Fig. 1d) in the pre-ageing regime (i.e., before structural transitions drive the condensate into a kinetically trapped state). The radius of gyration quantifies the overall degree of compactness of LCD proteins. Consistent with theoretical expectations and previous computational studies^60,73,74^, we observe a monotonic decrease in R*_g_* as the interface is approached, reflecting a gradual transition between different conformational states. LCDs within the condensate core adopt more extended conformations that maximize the intermolecular liquid network connectivity, whereas proteins in the dilute phase collapse into more compact conformations that maximize intra-LCD interactions, as reported in Fig. 1c. LCDs located across the interface display intermediate R*_g_* values, consistent with a gradual conformational crossover.

We next examine orientational order within the protein LCDs. The bond orientation (θ) is defined as the angle between the vector connecting two consecutive beads and the unit vector normal to the interface. This metric provides insight into chain alignment and is sensitive to short-range ordering and directional sequence correlations^75,76^. When analyzed as a function of distance from the condensate COM, our simulations (Fig. 1d) reveal no preferential bond orientation, as the mean value of θ remains close to 90*^?^* at every distance. Moreover, the distribution width, quantified by σ, is essentially constant across all positions, indicating the absence of enhanced local ordering of the LCD at the interface. Together, these observations suggest that interfacial effects on single-chain orientational order are minimal. Nevertheless, when comparing ageing kinetics for condensates with different surface areas—324 nm^2^ and 729 nm^2^—but identical total number of LCDs (Fig. 1e), we observe a markedly faster accumulation of inter-protein β-sheets in the system with larger surface to volume ratio. Figure 1e shows the time evolution of the normalized β-sheet concentration for both geometries, highlighting the speed up of condensate ageing with increasing interfacial area. These results independently confirm that the interface acts as a hotspot for structural transitions, consistent with the spatial probability profiles shown in Fig. 1b. Overall, while classical single-LCD properties such as conformational compactness and orientational order do not alone account for the preferential localization of structural transitions near the interface, the interface itself seems to play a decisive role in mediating condensate ageing. Therefore, a more comprehensive characterization is required to elucidate the mechanisms by which interfaces promote cross-β-sheet structure accumulation.

### C. Enhanced protein mobility and intermolecular rearrangements promote ageing at the condensate interface

To further elucidate the mechanisms that govern and promote ageing at the interface, we perform a more detailed characterization of both pre-ageing and post-ageing states. We first examine monomeric mobility, quantified through short-time diffusion coefficients (see Section SX of the Supplementary Material), which provide a measure of the dynamical state of the condensate. Figure 2a presents axial-cut spatial diffusion maps before and after ageing, alongside rendered images of spherical condensates pre- and post-incubation time. In the pre-ageing regime, monomers exhibit high mobility across all spatial regions. However, as condensates are incubated, LCD diffusion decreases markedly in areas where cross-β-sheet structures accumulate. While β-sheet formation is not the sole mechanism capable of reducing protein condensate mobility—density heterogeneities^77^, viscoelastic effects^59^, and other physicochemical processes^42,78^ also contributing—it remains the dominant driver of the observed diffusion slow down in our simulations. Notably, a pronounced low-diffusion region emerges at the interface, coinciding with enhanced interprotein β-sheet concentration. Beyond altering viscoelastic properties, reduced mobility at the interface may further limit molecular exchange between condensed and dilute phases^79,80^, a mechanism central to the functional dynamics of liquid-like physiological condensates^81^. Although our diffusion analysis provides valuable insight into how ageing modifies condensate material properties—particularly molecular mobility—it does not by itself explain why the interface actively promotes structural transitions over the bulk.

**Figure 2:**
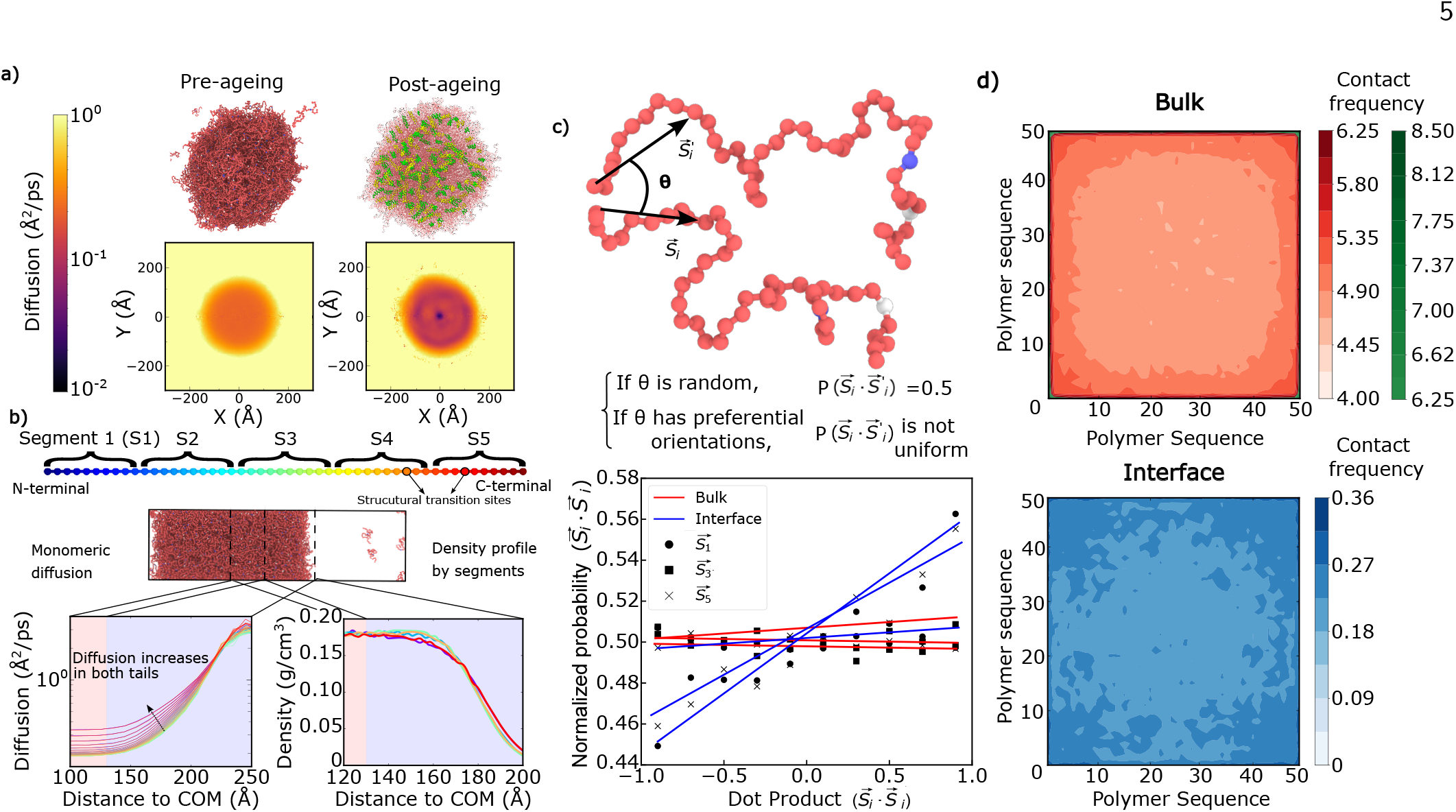
(a) Monomeric diffusion coefficient before (left) and after (right) ageing in spherical LCD condensates. The results are shown as projections onto the xy plane of the spherical droplet. Representative renders of the studied droplets are shown above the graphs. (b) Top: Scheme depicting the 10-residue long segments into which the sequence is classified. Middle: Render of a representative slab condensate. Bottom left: Monomeric diffusion as a function of distance from the condensate COM. Bottom right: Segment-resolved density profiles as a function of distance from the condensate COM. The plots follow the colour code shown in the top panel. (c) Normalized probability distribution of the dot product between end-to-end vectors of the same segment from different LCDs. Bulk and interface regions are considered separately, and segments S1, S3, and S5 are analysed (see panel (b) for segment definitions). A schematic representation of two LCD molecules is shown above the graph, defining the end-to-end vectors 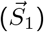 and the angle (θ). For random orientations, the normalized probability is uniform and approximately 0.5, while deviations indicate preferential intermolecular alignment. (d) Intermolecular contact frequency probability (in %) of the LCD for proteins located in the bulk and at the interface. The bulk contact map is shown using two colour scales due to the wide range of contact frequencies in this region.

Thus far, our analyses have focused on global system properties, as condensate density (Fig. 1b) or whole-chain observables, such as LCD dynamical conformation or mobility. However, to uncover the mechanisms primarily driving structural transitions at the interface, we perform a more detailed submolecular analysis. Consequently, we study monomeric mobility, measured as residue diffusion at short timescales (see Section SX of the SM), tracking the trajectory of each residue individually. In Fig. 2b (bottom left), we show the monomeric diffusion coefficient along the three directions of space as a function of the COM distance (results for the diffusion coefficient in the x-y plane and along the z direction, which yielded similar trends, are also reported in Section SX of the SM). The lines are colour-coded for each specific particle bead according to the scheme depicted in Fig. 2b (top). Interestingly, we find that terminal residues, at both ends of the LCDs, exhibit higher mobility regardless of their distance from the condensate COM. Moreover, monomeric diffusion increases for all residues as they approach the interface, with this enhancement becoming noticeable at approximately 150 °A from the condensate’s centre of mass. We identify two distinct effects: (i) the inherently higher mobility of the terminal segments, independent of the location across the condensate; and (ii) an interface-induced mobility enhancement (150–170 °A, see Fig. 2b). The latter establishes an interfacial regime in which increased mobility coincides with a local density sufficient to promote intermolecular interactions. Such interplay enhances the likelihood of forming intermolecular contacts at the interface and, consequently, the probability that high-density fluctuations trigger structural transitions into cross-β-sheets. We propose that this interplay—high mobility coexisting with a local density still sufficient to drive contacts—is precisely the physical origin of interfacial ageing. Therefore, our residue-level mobility calculations contribute to explain why the interface acts as a hotspot for condensate cross-β-sheet accumulation.

Furthermore, we analyse the condensate density profile by segmenting each protein LCD into five equal-length segments of 10 residues each. In Fig. 2b (bottom right), we show the density profile of each segment as a function of the COM distance. At the interface, (e.g., approximately 150 °A from the condensate’s COM) a statistically significant difference in local density between LCD segments emerges. The terminal segments (S1 and S5) exhibit a subtle density depletion prior to the overall density drop of the condensate, but a higher relative density compared to the central segments as we approach the dilute phase (e.g., in the 180– 200 °A range). This indicates that the terminal regions are preferentially located toward the interface, facing the dilute phase. In contrast, the central segments (S2, S3, and S4) maintain higher local densities at the interfacial region closer to the condensate core before undergoing a more abrupt density decay, leading to a crossover of the density probabilities within the interface. Altogether, these segmented density profiles suggest a preferential interfacial organization, in which terminal segments are enriched at the outer interface while central segments predominantly occupy a secondary coordination shell at the inner interface.

We further investigate this phenomenon by probing possible intermolecular alignment arising from the preferential arrangement of different protein segments at the interface. Specifically, we examine orientational correlations in the angle between end-to-end vectors 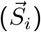 of pairs of segments belonging to different LCDs, as defined in Fig. 2c (top). This analysis is schematically illustrated in Fig. 2c, where we report the probability distribution of the dot product between normalized end-to-end vectors of segments from different protein replicas at the condensate. In the bulk (red curves in Fig. 2c), all segments exhibit a uniform distribution, consistent with random relative orientations (i.e., a probability centred around 0.5). In contrast, at the interface (blue curves in Fig. 2c), terminal LCD segments display a pronounced orientational preference, while central segments retain the uniform distribution characteristic of isotropic alignment in bulk conditions. These results demonstrate that intermolecular orientational order emerges selectively at the interface^82^ and is predominantly associated with terminal protein segments, where usually LARKS are located across the sequence^62^.

Next, we perform a detailed analysis of intermolecular protein interactions by computing contact frequency maps between all sequence residues. A contact is defined as effective—i.e., ensuring a significant enthalpic gain—when two residues are separated by less than 1.2 d, where d denotes the molecular diameter of each residue. In Fig. 2d, we compare two distinct regions of the condensate: the bulk (red and green maps) and the interface (blue map). In both regions, we observe a clear distinction between contacts involving terminal segments and those involving central segments, with terminal regions exhibiting consistently higher contact frequencies than the middle segments. As expected, the bulk region displays a larger overall number of contacts due to its higher local density compared to the interface. To quantify the degree of heterogeneity, we examine the ratio between the minimum and maximum contact frequencies at both near the interface and the core (120–140 °A and 30–50 °A distance range from the condensate’s COM, respectively), displaying a more pronounced differentiation between terminal and central segments contact probability at the interface. In principle, this enhanced contrast suggests that the spatial location of the structural transition sites could influence whether cross-β-sheet formation occurs preferentially at the interface. To directly assess the magnitude of this effect, we performed additional simulations in which the structural transition sites were repositioned at the chain centre (segment S3, Fig. 2b) and analysed the resulting ageing behaviour. As shown in Section SXI of the SM, the ageing rate still exhibits a maximum near the interface. Strikingly, relocating the transition-prone sites to the chain centre leaves the interfacial maximum unchanged (structural transition probability identical to Fig. 1b). This is a decisive test which demonstrates that interfacial ageing is governed by the physical organisation of the condensate, not by where the aggregation-prone motifs sit in the sequence.

Taken together, this residue- and segment-resolved analysis demonstrates that the interface is not merely a passive boundary, but a dynamically and structurally distinct region in condensates. Terminal segments exhibit enhanced mobility, preferential localization, and orientational ordering, whereas central segments remain comparatively uniform. These trends persist even when β-sheet-prone aggregation sites are relocated, indicating that interfacial ageing emerges from the intrinsic organisation and dynamics of the protein, rather than from the specific placement of the structural transition sites (e.g., LARKS in real protein sequences such as FUS, hnRNPA1, or TDP-43 among other numerous RNA-binding proteins across the human proteome22,40,53,62).

### D. Amphiphilic protein sequences amplify interfacial ageing

To bridge the gap between the minimal homopolymer LCD model and the architecture of real LCDs, we next introduce a more realistic protein model featuring chemically distinct regions across the sequence. As defined in Section II A, this amphiphilic LCD (A-LCD) consists of two interaction residue types, forming a more hydrophobic region with stronger intermolecular interactions, and a more hydrophilic region with weaker inter-protein interactions. A schematic representation of the A-LCD model is shown in Fig. 3a. After determining the critical solution temperature of this sequence (T*_c_* = 278 ± 2 K; see Section SII of the SM) and verifying that is almost identical to the original LCD sequence, we perform non-equilibrium condensate ageing simulations at T/T*_c_* = 0.92, as before. Analogously to the homopolymer LCD model (Fig. 1), we track the cross-β-sheet accumulation across the condensate space. These results are presented in Fig. 3b, where we compare the LCD and A-LCD models, together with the corresponding density profile for an A-LCD condensate in a slab geometry. Strikingly, structural transitions are more strongly enhanced toward the interface in the A-LCD compared to the LCD sequence, exhibiting a clearer maximum in the structural transition probability. This behaviour is also evident in Fig. 3a, which shows a rendering of a spherical A-LCD condensate alongside a sliced view, revealing a preferential localization of β-sheet fibrils at the interface. Importantly, increasing the asymmetry between the hydrophobic and hydrophilic interaction strength further amplifies the interfacial maximum in structural transition probability relative to the condensate bulk (see Fig. S9 in the SM).

**Figure 3:**
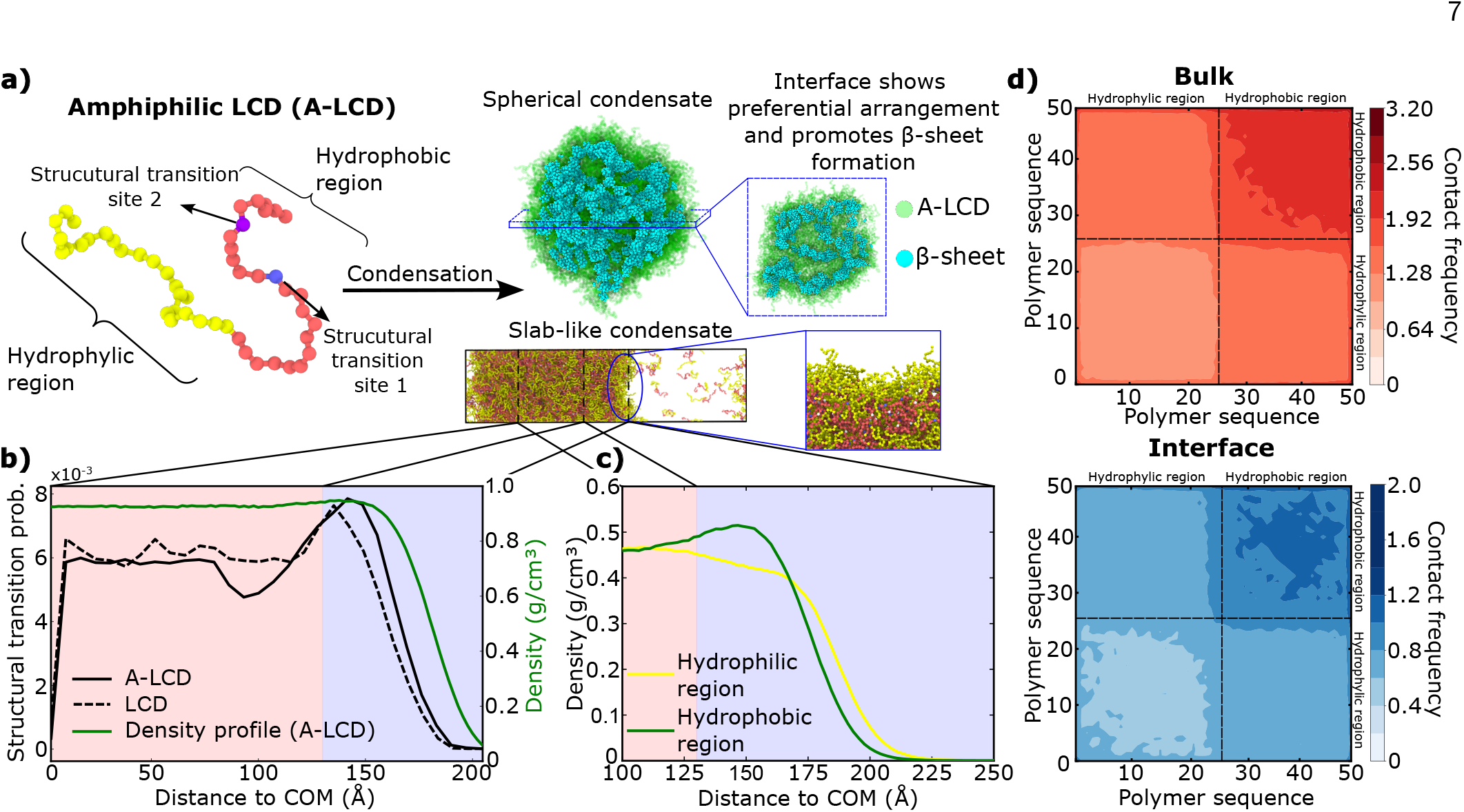
Amphiphilic proteins favour the appearance of β-sheet motifs at the condensate’s interface. (a) Schematic representation of the condensation process of the A-LCD into spherical and slab condensates. The sequence of the A-LCD is shown on the left, highlighting the structural transition sites and the hydrophobic and hydrophilic domains. A zoomed view of the slab-like condensate illustrates the preferential arrangement of A-LCD at the interface. (b) Structural transition probability of the A-LCD (solid black line) and homopolymer LCD (dotted black line), together with the density profile of an A-LCD condensate (green), as a function of distance from the condensate COM. (c) Density profiles of the hydrophilic and hydrophobic domains of the A-LCD sequence as a function of distance from the condensate COM. (d) Intermolecular contact frequency maps (in %) of A-LCDs in bulk (top) and at the interface (bottom). The two major domains of the A-LCD are indicated by dashed lines.

It is noteworthy to mention that the density profile in pre-ageing conditions, shown in Fig. 3b, exhibits a slight maximum located near the position of the peak of structural transition probability (post-ageing), a feature that is absent in the LCD density profile (Fig. 1b). To elucidate the origin of such behaviour, we decompose the density profile into contributions from the hydrophobic and hydrophilic domains. In the bulk (Fig. 3c), both regions display equivalent distributions. Upon approaching the interface, however, the two regions exhibit markedly distinct behaviours: the hy-drophobic domain develops a pronounced maximum on the inner side of the interface, while the hydrophilic region decreases monotonically and undergoes a sharp drop at the boundary of the dense phase, corresponding to the portion of the protein most exposed to the dilute phase to minimise the condensate surface tension. This observation is consistent with Fig. 3a, where the rendered configurations reveal a preferential localization of hydrophilic regions at the interface, while hydrophobic regions are displaced into a secondary coordination shell within the condensate. This spatial segregation promotes enhanced compaction through the association of hydrophobic regions, which experience stronger intermolecular interactions, thereby providing a mechanistic explanation for the emergence of a local density maximum near the interface.

To elucidate how this interfacial local density enhancement manifests at the molecular level, we examine the corresponding intermolecular contact maps of the A-LCD sequence. The contact frequency maps are shown in Fig. 3d, distinguishing between the bulk and interfacial regions of the condensate. A dashed line demarcates the hydrophobic and hydrophilic domains of the sequence in both maps. The maps reveal several key differences between the bulk and the interface. As expected, the highest contact frequencies occur within the hydrophobic region, particularly for hydrophobic–hydrophobic contacts, reflecting the stronger intermolecular interactions in this domain. In contrast, the hydrophilic region exhibits weaker and more homogeneous contact patterns, with both hydrophilic–hydrophilic and hydrophilic–hydrophobic interactions displaying lower and more uniform probabilities. Notably, the contrast between these regions becomes significantly amplified at the interface: the ratio of hydrophobic to hydrophilic-like contact frequencies increases to approximately 10:1 at the interface, compared to about 5:1 in the bulk. This enhanced disparity provides the molecular-level explanation for the strengthened interfacial structural cross-β-sheet transitions observed relative to the LCD system and directly arises from the asymmetric interaction patterning of the A-LCD sequence.

Overall, these results demonstrate that introducing chemical specificity does not substantially alter the global phase behaviour of the condensate, but it markedly influences the spatial organisation of ageing within the condensate. Compared to the minimal LCD model, the A-LCD sequence produces a pronounced enhancement of structural transitions near the interface, accompanied by a peak in the density profile, driven by preferential interactions among more hydrophobic segments. Crucially, this chemical asymmetry amplifies but does not create interfacial ageing—the effect is already present in the chemistry-free homopolymer—so sequence chemistry tunes the magnitude, not the location, of ageing. Taken together, these observations indicate that asymmetric patterns of chemical interactions along the sequence reinforce the role of the interface as a structurally and dynamically distinct region that promotes ageing.

### E. Surface tension minimization drives dynamic interfacial molecular ordering and ageing

Having established that chemical heterogeneity plays a major role in interfacial organisation and the accumulation of inter-protein structural transitions, we now turn to a more comprehensive study of pre-ageing condensate properties to further investigate the factors driving enhanced structural transitions at the interface. We begin by examining the monomeric mobility of A-LCD residues, quantified through short-time diffusion coefficients (see SM Section X), to assess how molecular motion varies as a function of distance from the condensate’s COM. These results are shown in Fig. 4a (bottom left), where each residue is represented by a separate curve. Compared to the homopolymer LCD (Fig. 2b), the symmetry between the two tails is lost: the hydrophilic tail exhibits higher diffusion than the hydrophobic tail. This behaviour is consistent with its lower binding network connectivity, as this region forms fewer intermolecular contacts (Fig. 2d), resulting in greater mobility. Nevertheless, this enhanced mobility alone does not explain the increased structural transitions at the interface, since the transition-prone sites are not located in this domain. In Figure S7 of the SM we present a two-dimensional analysis of monomeric mobility before and after ageing, highlighting the increased mobility at the condensate interface and its subsequent arrest following the formation of inter-protein β-sheets through the hydrophobic domain.

**Figure 4:**
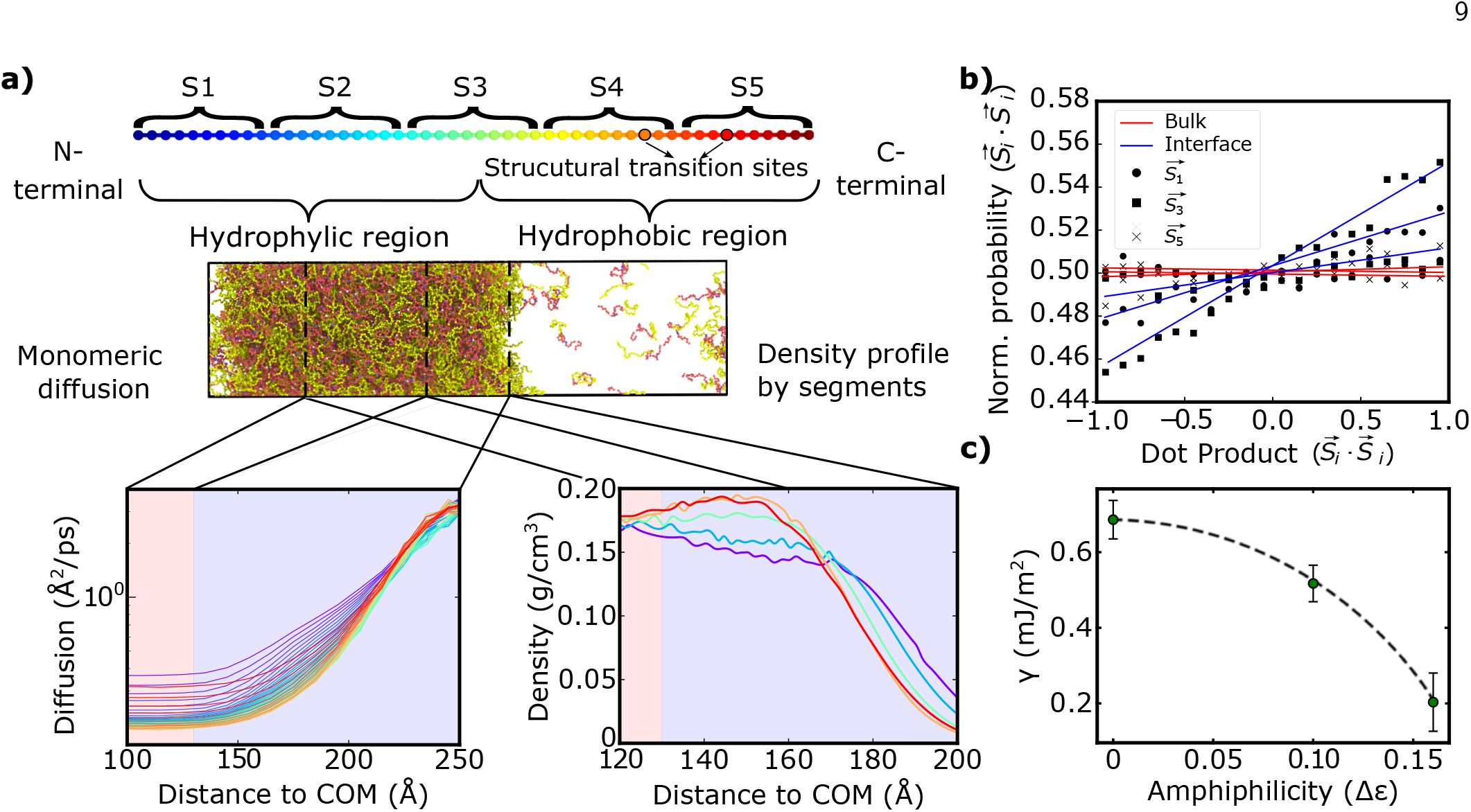
(a) Top: Schematic depiction of the A-LCD sequence, showing the colour code, structural transition sites, and different segments considered in our analysis. Middle: Rendered image of a representative slab simulation. Bottom left: Monomeric diffusion as a function of distance from the condensate COM. Bottom right: Segment-resolved density profiles versus distance from the condensate COM. (b) Normalized probability of the dot product between end-to-end vectors of the same segment from different LCDs. Bulk (red) and interface (blue) regions are considered separately, analysing segments S1, S3, and S5 (see panel (a) for segment definitions). Distances of 50–60 °A and 120–130 °A from the condensate COM define the bulk and interfacial regions, respectively. (c) Surface tension (γ) as a function of the amphiphilicity degree expressed in terms of the used ε value in the Wang–Frenkel potential used for describing the hydrophobic vs. hydrophilic domain interaction strength. Error bars represent standard deviation.

Besides monomeric diffusion, we also examine the condensate density profile per segment (Fig. 4a, top). The resulting profiles (Fig. 4a, bottom right) reveal a striking spatial organisation within the condensate. In the bulk (not shown), all segments exhibit similar densities, consistent with a homogeneous interior. However, approaching the interface, the hydrophobic and hydrophilic segments display the same behaviour observed in Fig. 3c for the respective two major protein domains. Hydrophobic segments occupy the inner side of the interface (*∼*150 °A), whereas more hydrophilic segments face the dilute phase at greater distances (>180 °A). Next, we assess whether this interfacial ordering induces intermolecular alignment by measuring the orientation of each segment using the parameter θ, as defined in Section II C. Figure 4b shows the results for the bulk (50–60 °A from the condensate COM) and within interfacial regions (120–130 °A). In the bulk (red curves), all segments exhibit uniform orientation, with values centred around 0.5, reflecting the absence of directional ordering. In contrast, at the interface (blue curves), central segments display a pronounced preferential orientation, while terminal segments show a weaker, yet significant, orientational bias. We attribute the stronger orientation of central segments to their mixed hydrophobic–hydrophilic composition and their role in mediating the interfacial arrangement: aligning the hydrophobic tails at the inner interface while maintaining the hydrophilic domains at the outer layer. Overall, this analysis reveals a spatial organisation distinct from the homopolymer LCD, where the chemical nature of the segments plays a more important role than their specific position along the sequence.

Lastly, we investigate how protein intermolecular arrangements at the interface—which ultimately promote structural transitions (Fig. 3)—arise from the condensate’s propensity to minimise its surface tension. To test this point, we compute the interfacial free energy of pre-aged condensates (details in Section SXII of the SM) as a function of increasing protein amphiphilicity. We define the degree of amphiphilicity as the difference between the interaction strength of the hydrophobic and hydrophilic domains, expressed in terms of ε in the Wang–Frenkel potential (see Section SI of the SM). As shown in Fig. 4c, the surface tension decreases monotonically with increasing chemical sequence asymmetric interactions. This behaviour is fully consistent with an amphiphilic organisation at the interface (Figs. 3a and 4a), where the preferential orientation and clustering of chemically alike regions enable the hydrophilic domains to act as surfactants, stabilizing the interface and lowering its associated free-energy cost, in line with recent findings that sequence composition and blocky patterning modulate condensate interfacial properties^83–86^.

Taken together, these results show that introducing chemical heterogeneities within the sequence leads to a condensate spatial organisation distinct from that of the homopolymer LCD, further enhancing its propensity to undergo β-sheet structural transitions at the interface (Fig. 3b). The mobility profiles, segment-resolved density distributions, and orientational correlations consistently indicate that the behaviour of the A-LCD is dominated by the chemical nature of its domains rather than merely on their position along the sequence. Such organisation is driven by the condensate’s tendency to minimise its surface tension, which exhibits a systematic decrease with increasing chemical asymmetry without barely affecting condensate stability (Fig. S1 shows the phase diagrams of the different LCD sequences). The resulting interface, stabilized yet dynamic and chemically organized, provides a favourable environment for the emergence and localization of ageing-related structural transitions.

## III. CONCLUSIONS

This work identifies the physical origin of ageing at biomolecular condensate interfaces. Using coarse-grained molecular dynamics simulations of minimal protein models capable of both phase separation and inter-protein β-sheet formation, we show that the condensate interface is not merely a boundary between dense and dilute phases but a distinct physical environment that promotes structural transitions. Across all systems studied, ageing emerges preferentially at the interface through a common mechanism that does not depend on the specific chemistry of a given protein sequence.

As summarised in Fig.5, unlike the condensate interior, the interface uniquely combines enhanced molecular mobility, local density fluctuations and inter-protein alignment. Proteins exhibit increased mobility at the condensate surface, particularly at terminal segments, which facilitates encounters between regions capable of undergoing structural transitions (Fig.5, top left). At the same time, the interface induces a dynamic inter-protein orientational order absent from the bulk. That is, at the interface, proteins remain mobile and continuously rearrange, yet display a preferential alignment of terminal domains rather than the largely isotropic organisation observed within the condensate interior (Fig. 5, top right). The coexistence of mobility, alignment and local concentration creates ideal conditions for the nucleation of inter-protein β-sheet structures.

**Figure 5:**
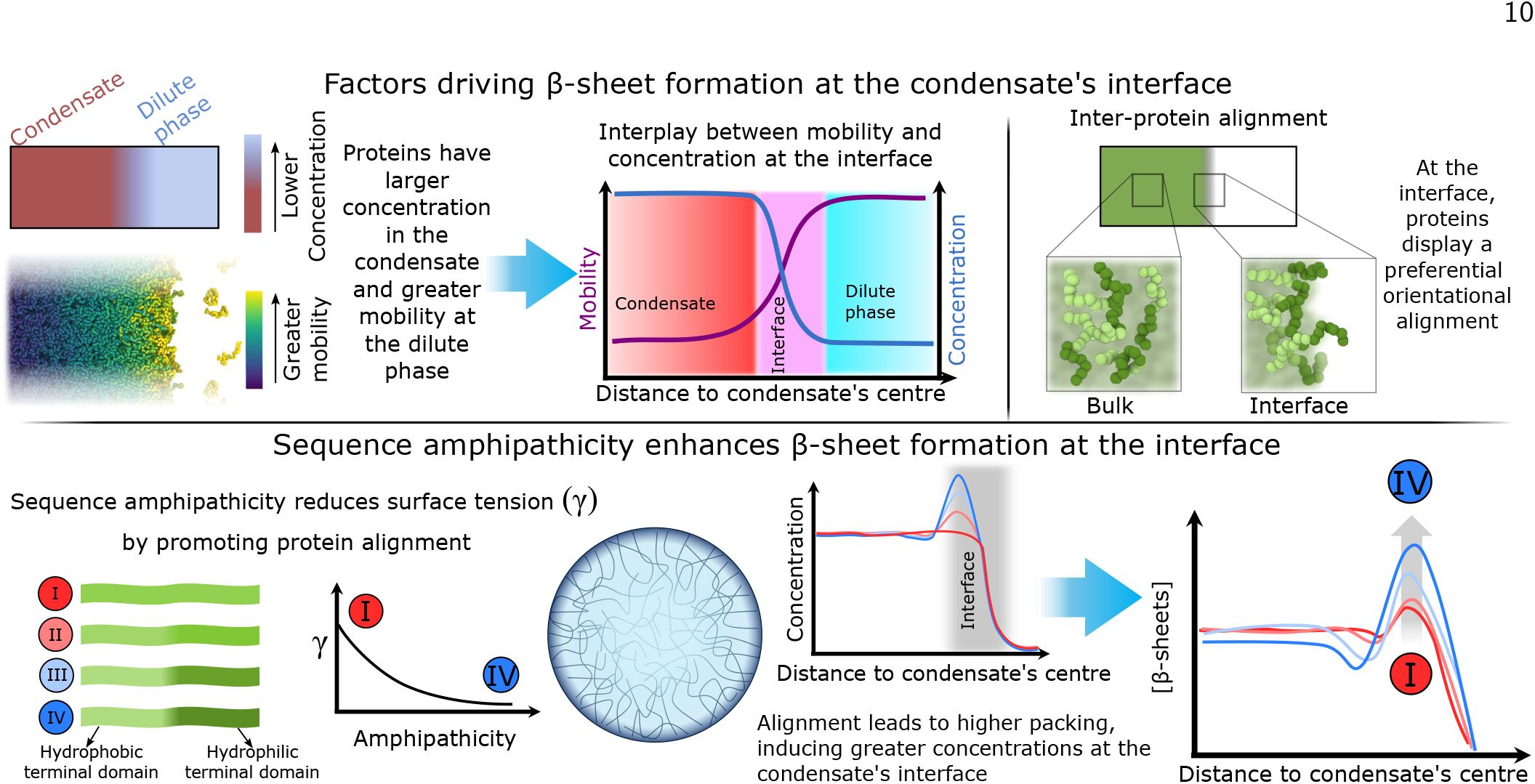
Top: Schematic summary of the factors contributing to the enhanced formation of inter-protein β-sheets at the condensate interface. Bottom: Schematic representation of the impact of sequence amphipathicity in condensate inter-protein β-sheet formation. The plots are illustrative.

Sequence asymmetry strengthens this generic interfacial mechanism. In amphiphilic proteins, hydrophilic domains preferentially orient towards the dilute phase, while hydrophobic regions remain buried beneath the interface. This surfactant-like organisation lowers the interfacial free energy and reinforces the inter-protein orientational alignment already favoured by the interface. As a result, proteins pack more efficiently near the interface, generating local density maxima and concentration fluctuations that further promote cross-β-sheet nucleation. Thus, amphiphilicity amplifies interface-driven ageing by tuning surface tension, packing and molecular alignment, but it is not required for ageing to begin at the interface. Sequence chemistry therefore modulates the magnitude of interface-driven ageing but does not determine its origin.

Several important questions remain open. Future work should investigate how these mechanisms operate in longer and more complex proteins, including multidomain systems containing folded domains; how proximity to criticality influences interfacial ordering through changes in surface tension; and how explicit solvent effects contribute to interfacial ageing and to the formation of hollow condensates observed experimentally. Addressing these questions will help establish how universal the physical principles identified here remain across the broad spectrum of biological condensates.

Taken together, our results suggest that interface-driven ageing in biomolecular condensates is governed by generic physical features of the interface rather than by sequence-specific chemistry alone. The coexistence of mobility, alignment and local concentration at condensate interfaces provides a general physical framework for understanding why diverse condensate-forming proteins, including FUS, hn-RNPA1 and α-synuclein, exhibit preferential ageing at interfaces. This shifts attention from the chemistry of individual proteins to the physical properties of the interface itself, highlighting surface tension, interfacial composition and dynamic molecular ordering as potential control parameters for delaying pathological liquid-to-solid transitions.

## IV. DATA AVAILABILITY

The data that supports the findings of this study are available within the article and its Supplementary Material.

## V. CODE AVAILABILITY

The necessary configuration and LAMMPS script to run a non-equilibrium ageing simulation can be found in the repository GitHub link for the repository (https://github.com/Alex-castro-quim/Interface)

## VI. ACKNOWLEDGEMENTS

I. S.-B. acknowledges funding from Derek Brewer scholarship of Emmanuel College and EPSRC Doctoral Training Programme studentship, number EP/T517847/1, Ramon y Cajal fellowship (awarded to J.R.E.), as well as the UKRI EPSRC under the UK Government’s guarantee scheme (EP/Z002028/1), following successful evaluation by the ERC (Consolidator Grant awarded to R.C.G.) under the European Union’s Horizon Europe research and innovation programme. A. R. T. acknowledges funding from the European Union Horizon 2020 research and innovation programme (grant agreement 803326 to R.C.-G.). J. R. E. acknowledges funding from the Ramon y Cajal fellowship (RYC2021-030937-I), the Spanish scientific plan and committee for research reference PID2022-136919NA-C33, and the European Research Council (ERC) under the under the European Union’s Horizon Europe research and innovation program (grant agreement no. 101160499). C. and J. L. -M. acknowledge funding from the ERC grant agreement no. 101160499 (awarded to J. R. E). This work has been performed using resources provided by Archer2 (https://www.archer2.ac.uk/) funded by EPSRC Tier-2 capital grant EP/P020259/e829. The authors also thankfully acknowledge RES computational resources provided by Mare Nostrum 5 through the activity 2024-3-0001.

## VII. AUTHORS CONTRIBUTIONS STATEMENT

I.S-B. R.C.G., and J.R.E conceived the project, and together with A.C., A.R.T., J.L-M., M.P, P.A. and A.O. carried it out the simulations and data analysis; J.R.E and I.S-B. supervised the project and J.R.E and R.C-G. provided computational means and funding. A.C. wrote the paper and all authors contributed equally to revising it.

