## Supplementary Material for "The physical origin of ageing at biomolecular condensate interfaces"

### Supplementary Material: The physical origin of ageing at biomolecular condensate interfaces

Alejandro Castro[1], Juan Luengo-Márquez[1], Andres R. Tejedor[1,2], Rosana Collepardo-Guevara[2,3,4,5], Marcell Papp[6], Paolo Arosio[6], Alberto Ocaña[5,7,8], Ignacio Sanchez-Burgos[2,3*] and Jorge R. Espinosa[1,2,3,5,8*]
*[1] Department of Physical-Chemistry, Universidad Complutense de Madrid, Av. Complutense s/n, 28040, Madrid, Spain.*
*[2] Yusuf Hamied Department of Chemistry, University of Cambridge, Lensfield Road, Cambridge CB2 1EW, United Kingdom*
*[3] Maxwell Centre, Cavendish Laboratory, Department of Physics, University of Cambridge, J J Thomson Avenue, Cambridge CB3 0HE, United Kingdom.*
*[4] Department of Genetics, University of Cambridge, Cambridge CB2 3EH, United Kingdom.*
*[5] PhAsIca Biosciences S.L, Calle Velázquez, 27, 28001 Madrid, Spain*
*[6] Institute of Biochemistry, Department of Biology, ETH Zurich, Zurich, 8093, Switzerland*
*[7] Experimental Therapeutics in Cancer Unit, Instituto de Investigación Sanitaria San Carlos (IdISSC), and CIBERONC, Madrid, Spain*
*[8] Multidisciplinary Institute, Complutense University of Madrid, Paseo Juan XXIII, 1, Madrid 28040, Spain*
(Dated: 12th June 2026)

#### SI. MODEL AND METHODS

In our simulations, we represent the proteins as coarse-grained chains with one bead per monomer. The potential energy of the coarse-grained force field is defined as follows:

$$E = E_{\text{Bonds}} + E_{\text{Hydrophobic}} \ . \tag{S1}$$

Bonded interactions are modelled by a harmonic potential

$$E_{\text{Bonds}} = \sum k(r_i - r_0)^2, \tag{S2}$$

where $k = 9.6\text{kJ/mol} \cdot \text{Å}^2$. The equilibrium bond length is $r_0 = 3.81$ Å between bonded monomers.

Table S1: Comparison of interaction parameters across models

| Type 1 | Type 2 | Model | | | | | |
|---|---|---|---|---|---|---|---|
| | | LCD | | Amphiphilic LCD (I) | | Amphiphilic LCD (II) | |
| | | $\epsilon$ | $\sigma$ | $\epsilon$ | $\sigma$ | $\epsilon$ | $\sigma$ |
| * | * | 0.50 | 6.00 | 0.55 | 6.00 | 0.58 | 6.00 |
| 3 | 3 | 4.00 | 6.00 | 4.00 | 6.00 | 4.00 | 6.00 |
| 5 | 5 | 4.00 | 6.00 | 4.00 | 6.00 | 4.00 | 6.00 |
| 9 | 1 | – | – | 0.50 | 6.00 | 0.50 | 6.00 |
| 9 | 9 | – | – | 0.45 | 6.00 | 0.42 | 6.00 |
| 9 | 2 | – | – | 0.50 | 6.00 | 0.50 | 6.00 |
| 9 | 4 | – | – | 0.50 | 6.00 | 0.50 | 6.00 |
| 9 | 3 | – | – | 0.50 | 6.00 | 0.50 | 6.00 |
| 9 | 5 | – | – | 0.50 | 6.00 | 0.50 | 6.00 |
| 9 | 6 | – | – | 0.50 | 6.00 | 0.50 | 6.00 |
| 9 | 7 | – | – | 0.50 | 6.00 | 0.50 | 6.00 |

Hydrophobic interactions are implemented via the Wang-Frenkel potential

$$E_{\text{Hydrophobic}} = \sum_i \sum_{j<i} \epsilon_{ij} \alpha \left( \left[ \frac{\sigma_{ij}}{r} \right]^{2\mu} - 1 \right) \left( \left[ \frac{r_c}{r} \right]^{2\mu} - 1 \right)^{2\nu_{ij}} \ , \tag{S3}$$

being

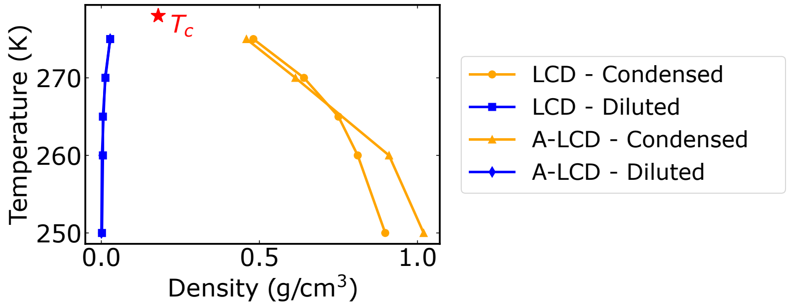


Figure S1: Phase diagrams for (a) homopolymer model (b) Amphiphilic polymer model, calculated as described in Section SII. We also provide the linear fit used for equations S5 and S6 as dashed lines.

$$\alpha = 2\nu \left(\frac{r_c}{\sigma_{ij}}\right)^{2\mu} \left[\frac{1+2\nu_{ij}}{2\nu_{ij}\left(\left(\frac{r_c}{\sigma_{ij}}\right)^{2\mu}-1\right)}\right]^{2\nu_{ij}+1}, \tag{S4}$$

where the excluded volume of the different residues is given by $\sigma_{ij}$, $r$ is the distance between the $ij$ particles, $\epsilon_{ij}$ is the energy interaction parameter, $r_c = 3\sigma_{ij}$ is the cut-off of the potential between those beads and $\mu$=1 and $\nu_{ij}$ control the functional shape of the potential. The values of $\sigma_{ij}$ and $\epsilon_{ij}$ are shown in Table S1. We note that atom types 3 and 5 are reserved for the structural transitions sites post-ageing, as defined in the main text. Atom type 9 is reserved for the hydrophilic half in the A-LCD sequences, while the remaining ones define the hydrophobic part of the sequence. The table shows the interactions in LAMMPS style, where a new definition of any pairwise interaction overrides the previously defined one.

#### SII. PHASE DIAGRAMS

As outlined in the main text, we compute the critical temperature for phase separation for the different models through NVT simulations via the Direct Coexistence method. Here, we perform simulations below the critical temperature, and measure the density of the coexisting condensed and diluted phases. When the phase diagram is calculated via this method, the critical point of the phase diagrams is estimated using the universal scaling law of coexistence densities near a critical point [1]

$$(\rho_l(T) - \rho_v(T))^{3.06} = d\left(1 - \frac{T}{T_c}\right), \tag{S5}$$

and the law of rectilinear diameters [2]

$$\frac{\rho_l(T) + \rho_v(T)}{2} = \rho_c + s_2(T_c - T), \tag{S6}$$

where $\rho_l$ and $\rho_v$ refer to the coexisting densities of the condensed and diluted phases respectively, $\rho_c$ is the critical density, $T_c$ is the critical temperature, and $d$ and $s_2$ are fitting parameters.

The phase diagrams are shown in Fig. S1, where both, homopolymer and Amphiphilic system are represented. The parameters obtained from this result are shown in table S2

#### SIII. AGEING ALGORITHM

We perform our ageing simulations allowing the formation of inter-protein β-sheets to take place, according to the scheme developed in Refs. [3–5]. We first define two beads within the 50-bead polymers (beads 38 and 44 in the

Table S2: Phase diagram parameters

| | Homopolymer model | Amphiphilic model |
|---|---|---|
| $T_c\ (K)$ | 277.92 | 277.78 |
| $\rho_c\ (g/cm^3)$ | 8.27 | 7.94 |

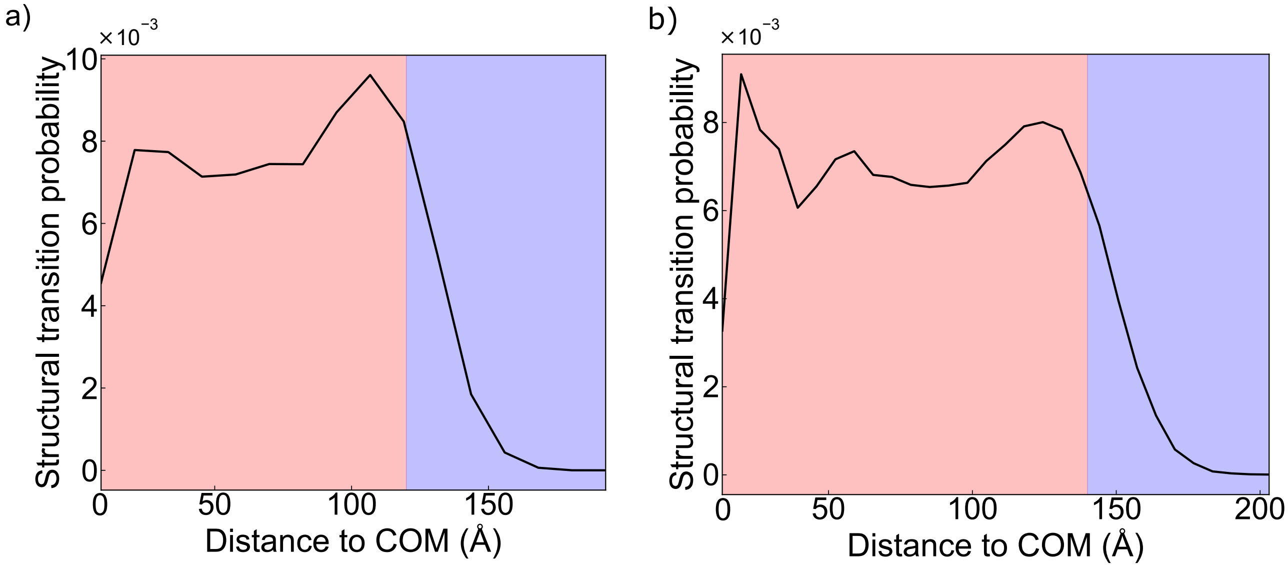


Figure S2: β-sheet formation probability as a function of the distance to the centre of mass of the condensate (COM) of spherical systems containing (a) ∼1000 LCD replicas and (b) ∼1500 LCD replicas. The red shaded region corresponds to the bulk, while the blue shaded region denotes the interface.

sequence) that are prone to forming inter-protein secondary structure. These regions mimic the low-complexity aromatic-rich kinked segments (LARKS). Our ageing simulations consist of non-equilibrium Molecular Dynamics runs in which, when the aforementioned regions come into contact satisfying a geometric criterium, the pairwise interaction energy between the associated residues changes. The geometric criterium requires that: (i) at least four LARKS of different proteins come into contact simulataneously, and (ii) these lie under a cutoff distance (9.75 Å) relative to each other, that is set as to observe the formation of inter-protein β-sheets within an accessible timescale. This modification is implemented in LAMMPS by means of the *fix bond/react* command[6].

We perform the ageing simulations in the canonical ensemble. Example files of this algorithm can be found in the online repository (GitHub link for the repository ( https://github.com/Alex-castro-quim/Interface), where we have included configuration files for all polymers, as well as the LAMMPS scripts and associated files needed to run a non-equilibrium simulation.

With our computational setup, we are capable of performing simulations at a speed of 600 timesteps per CPU second, which allows us to simulate approximately 200 ns per day for systems of ∼50000 particles (i.e., ∼ 1000 protein replicas) and 112 CPUs per simulation (tested with Intel Xeon Platinum 8160 CPU).

#### SIV. SPHERICAL STRUCTURAL TRANSITION PROBABILITY LOCATION

As stated in the main text, here we provide the β-sheet formation location results for the spherical condensates. In Fig. S2a and Fig. S2b we find results for both systems.

In spherical geometries, the population of particles within each radial shell is not uniform but instead varies systematically with the distance to the centre of the condensate due to purely geometric volume effects. Specifically, the volume associated with a spherical shell increases with radius, leading to a larger number of polymers or monomers being sampled at larger distances even in the absence of any underlying physical heterogeneity. This fact explains the maximum observed in both systems at around 10 Å, as the volume effect overestimates the transition probability.

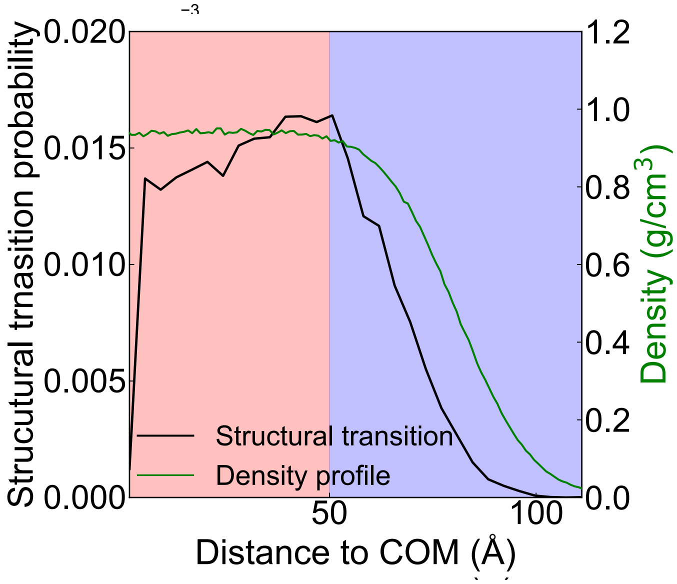


Figure S3: β-sheet formation probability as a function of the distance to the centre of mass (COM) of the LCD system condensate for a slab system with ∼700 nm$^2$. The red shaded region corresponds to the bulk, while the blue shaded region denotes the interface. Along with these areas, the density profile is plotted as a green line.

#### SV. INTERFACIAL MAGNITUDE STRUCTURAL TRANSITION PROBABILITY LOCATION

As mentioned in the main text, we considered two systems with different surface area. In this section we show the results for the structural transition probability in a system for which interfacial is ∼700 nm$^2$ in Fig. S3.

Note that this system has the same volume and number of particles as the other slab system (∼300 nm$^2$), and thus the longitude of the condensate is drastically reduced. This effect translates in a narrow bulk region, spanning only around 100 Å, and implying that interfacial effects dominate a much larger fraction of the condensate volume. As a result, the maximum in this system is less pronounced since there is more overlap between bulk and interface regions. This overlap smears out spatial distinctions, reducing the contrast of the interfacial maximum.

#### SVI. NUCLEATION

We distinguish between nucleation events—corresponding to the initial formation of a stable β-sheet seed—and structural transition events associated with elongation. In Fig. S4, we explicitly analyse the spatial distribution of nucleation events to determine where they preferentially occur within the condensate. The resulting profiles follow the same qualitative trend observed for elongation, an enhanced occurrence near the interface. However, due to the intrinsically low frequency of nucleation events, the statistical sampling is significantly poorer than for elongation, leading to larger fluctuations in the spatial distributions. Despite this reduced statistical sampling, the consistency in trends supports the same physical picture discussed in the main text.

#### SVII. AGEING CURVES

We used 10 independent random seeds to account for the stochastic variability of the phenomena studied. In Fig. S5 we show the relative β-sheet concentration as a function of the time for the two slab systems, whose comparison can be found in Fig. 1e of the main text. We plot the different curves for individual seeds (coloured lines) and the average of those lines (black line).

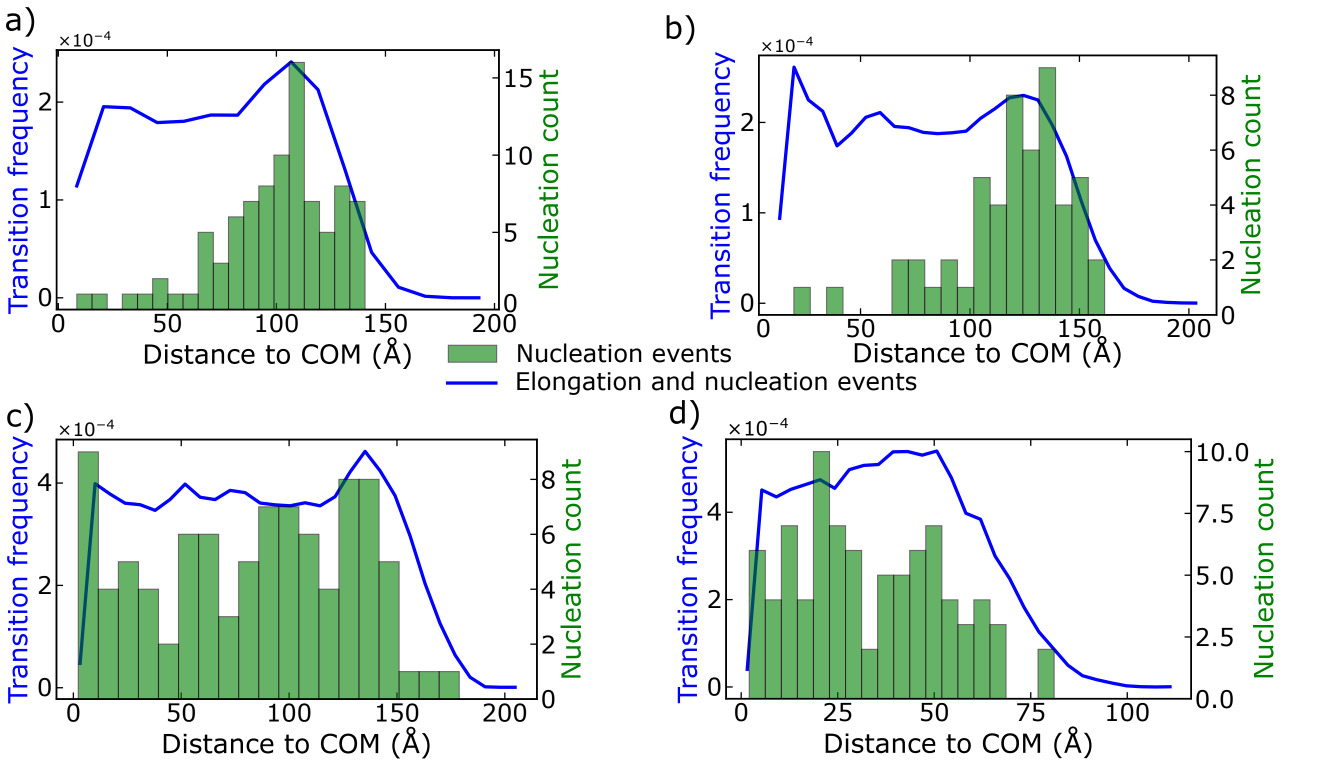


Figure S4: Nucleation and β-sheet formation probability as a function of the distance to the centre of mass of the biggest cluster (COM) of (a) spherical systems containing ∼1000 polymer replicas, (b) spherical systems containing ∼1500 replicas, (c) slab system with ∼300 nm$^2$ interfacial area and (d) slab system with ∼700 nm$^2$. Nucleation events are represented as green bars and correspond to the raw nucleation counts, whereas the transition frequency (blue lines) is normalized by the local volume. All results correspond to the LCD model

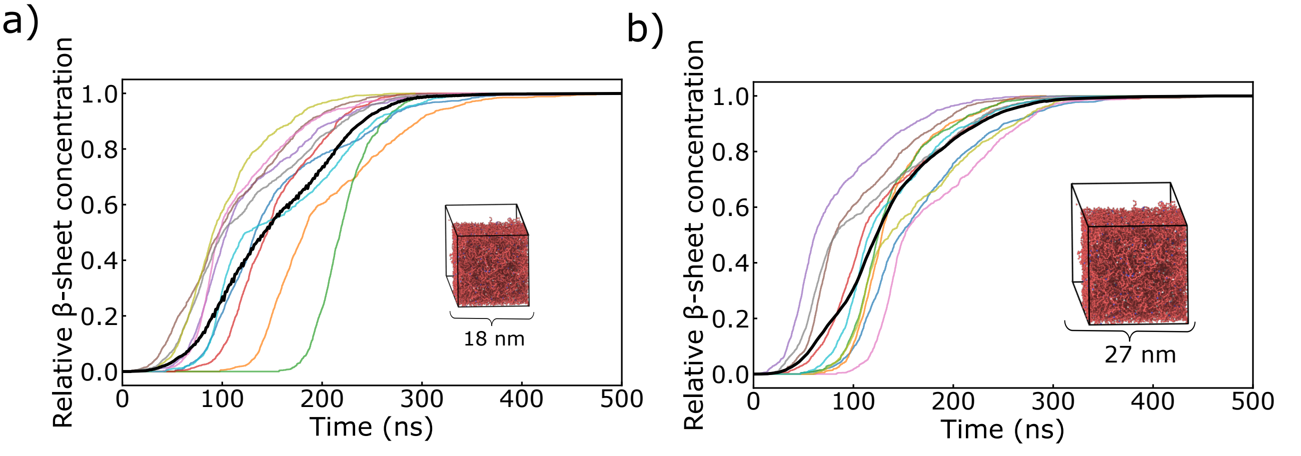


Figure S5: Relative β-sheet concentration for the LCD system as a function of time of slab systems with (a) ∼300 nm$^2$ interfacial area and (b) ∼700 nm$^2$. Each seed is represented as a unique curve and the average is plotted as the black curve.

#### SVIII. FREQUENCY CONTACT MAPS

For this analysis, we consider an effective contact to take place when two residues are found at a distance below $1.2\sigma_{ij}$, being $\sigma_{ij}$ the average molecular diameter of the two residues $i$ and $j$, and 1.2 a slightly larger distance to the minimum in the potential ($\sim 1.12\sigma_{ij}$) between the $i$ and $j$ monomers.

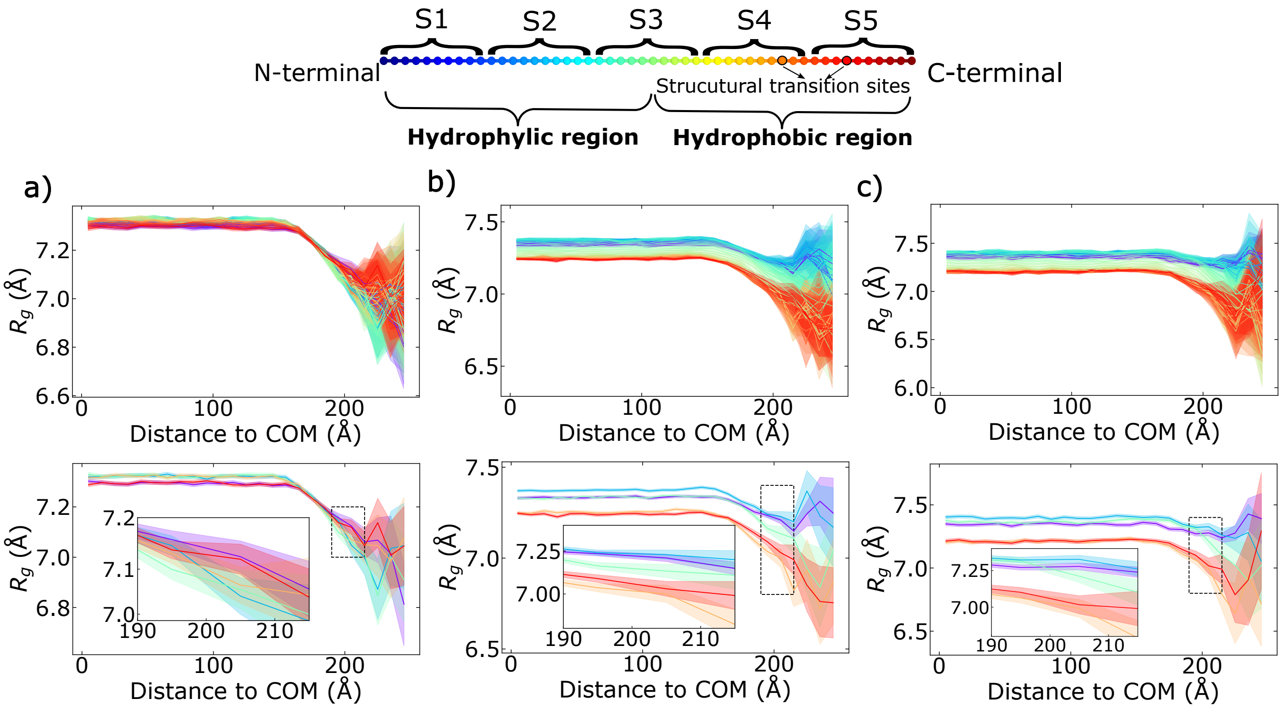


Figure S6: Top: Scheme depicting the 10-bead long segments that we clasiffy our sequence into. Bottom: Radius of gyration as a function of the distance to the centre of mass of the condensate, showing only S1, S2, S3, S4, S5 as indicated in the sheme above, of (a) LCD model (b) Amphiphilic LCD I model and (c) Amphiphilic LCD II model. The error is represented by a shaded outline for each curve.

#### SIX. RADIUS OF GYRATION BY SEGMENTS

In Fig. S6, we show the radius of gyration by segments for the three models we have designed. Here the lines are coded for each specific segment according to the scheme in Figure S6 (top). In the top of every panel we show every possible 10 bead segment plotted, while at the bottom we find 10 equispaced segments corresponding to S1, S2, S3, S4 and S5. The error of each curve is represented as a shaded outline of the same colour as the curve.

The LCD model (Fig. S6a) shows similar and nearly constant $R_g$ in the bulk, indicating a homogenous conformational state away from the interface. Nonetheless, subtle differences are observed between different segments: the tail segments exhibit a consistent smaller radius of gyration in the bulk compared to middle segments. We attribute this behaviour to an enhanced intermolecular connectivity of the tail beads in bulk, which allows them to explore a broader conformational landscape and thus being able to adapt a more compact configurations. Upon approaching and crossing the interface, a systematic decrease in $R_g$ is observed for all segments. During this decrease a crossover between tail and middle segments emerges. After the crossover region, tail segments display a maximum in this property whereas middle segments exhibit a minimum. This behaviour is consistent with previous observations reported by Wang *et al.*[7] who also described segment-specific conformational rearrangements near interfaces.

Both amphiphilic models (Fig. S6b,c) display very similar characteristics, and, in contrast to the LCD model, exhibit a more segment-dependent behaviour (a bigger contrast is found in the Amphiphilic II model). Once again, in bulk, segments show similar and nearly constant $R_g$ values. Throughout all distances, hydrophobic regions show more compact configurations than the hydrophilic segments. The crossover behaviour observed in the homopolymer is thus replaced by a chemically driven differentiation, where conformational rearrangements at the interface are primarily governed by the nature of the segment rather than by its position along the chain. This highlights that, in the Amphiphilic models, interfacial conformational changes are dictated by amphiphilic organization rather than purely geometric effects.

#### SX. DIFFUSION ANALYSIS METHODOLOGY

The diffusion analysis was performed using post-processing of molecular dynamics (MD) trajectories. The objective is to determine the short-time diffusion coefficient of individual beads within polymer chains using the relation between mean squared displacement (MSD) and diffusion. Once again, these quantities should be interpreted as short-time, frame-to-frame mobility estimates derived from finite-time displacements, rather than true long-time diffusion coefficients

The diffusion coefficient is calculated using the relation

$$\langle r^2(t)\rangle = 2dDt, \tag{S7}$$

where $d$ is the dimensionality of the system.

Accordingly, we consider the instantaneous diffusion coefficients defined as

$$D_{xyz} = \lim_{\Delta t \to 0} \frac{\Delta r^2}{6\Delta t}, \tag{S8}$$

$$D_{xy} = \lim_{\Delta t \to 0} \frac{\Delta r_{xy}^2}{4\Delta t}, \tag{S9}$$

$$D_{z} = \lim_{\Delta t \to 0} \frac{\Delta r_{z}^2}{2\Delta t}. \tag{S10}$$

In practice we compute the instantaneous diffusion coefficients in eq. S8, S9 and S10 by taking the limit as the time between two consecutive frames, and computing the square displacement between those frames.

Here:

- $D_{xyz}$ corresponds to three-dimensional diffusion ($d = 3$),
- $D_{xy}$ corresponds to in-plane diffusion ($d = 2$),
- $D_z$ corresponds to one-dimensional diffusion along $z$ ($d = 1$).

In this section we also show the results for $D_{xy}$ and $D_z$ for the LCD model and A-LCD I model. These results can be found in Fig. S7a,b respectively. The discussion to these results is analogue to the one done in the main (Sections C and E in results).

#### SXI. MIDDLE LARK

As mentioned in the main text, in Fig. S8b we show the comparison between the LCD-tail and the LCD-middle. The main difference between these homopolymers is the location of the structural transition sites, going from positions 44 and 38 for the LCD-tail to 22 and 28 for the LCD-middle.

The structural transition probabilities are identical, indicating that the greater probability of tail-tail interactions at the interface plays a negligible role in promoting ageing at the condensate interface.

#### SXII. SURFACE TENSION CALCULATION AND ERROR

The surface tension was computed using the mechanical (pressure tensor) definition for a planar interface normal to the $z$-direction:

$$\gamma = \frac{L_z}{2}\left(P_{zz} - \frac{P_{xx} + P_{yy}}{2}\right), \tag{S11}$$

where $L_z$ is the box length in the elongated direction, perpendicular to the interface, and $P_{xx}, P_{yy}, P_{zz}$ are the diagonal components of the pressure tensor. The factor $1/2$ accounts for the presence of two interfaces in the slab geometry.

The instantaneous surface tension at timestep $t$ is therefore:

$$\gamma(t) = \frac{L_z}{2}\left(P_{zz}(t) - \frac{P_{xx}(t) + P_{yy}(t)}{2}\right). \tag{S12}$$

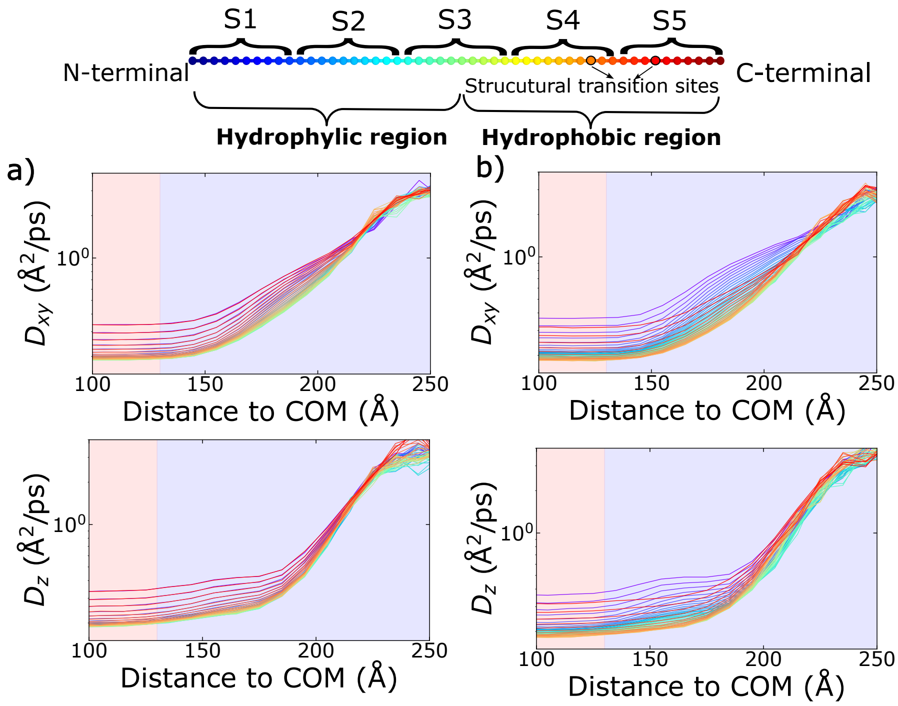


Figure S7: Top: Scheme depicting the 10-bead long segments that we clasiffy our sequence into. Bottom: Difussion as a function of the distance to the centre of mass of the biggest cluster (COM) of (a) homopolymer model and (b) Amphiphilic I model. The error is represented by a shaded outline for each curve.

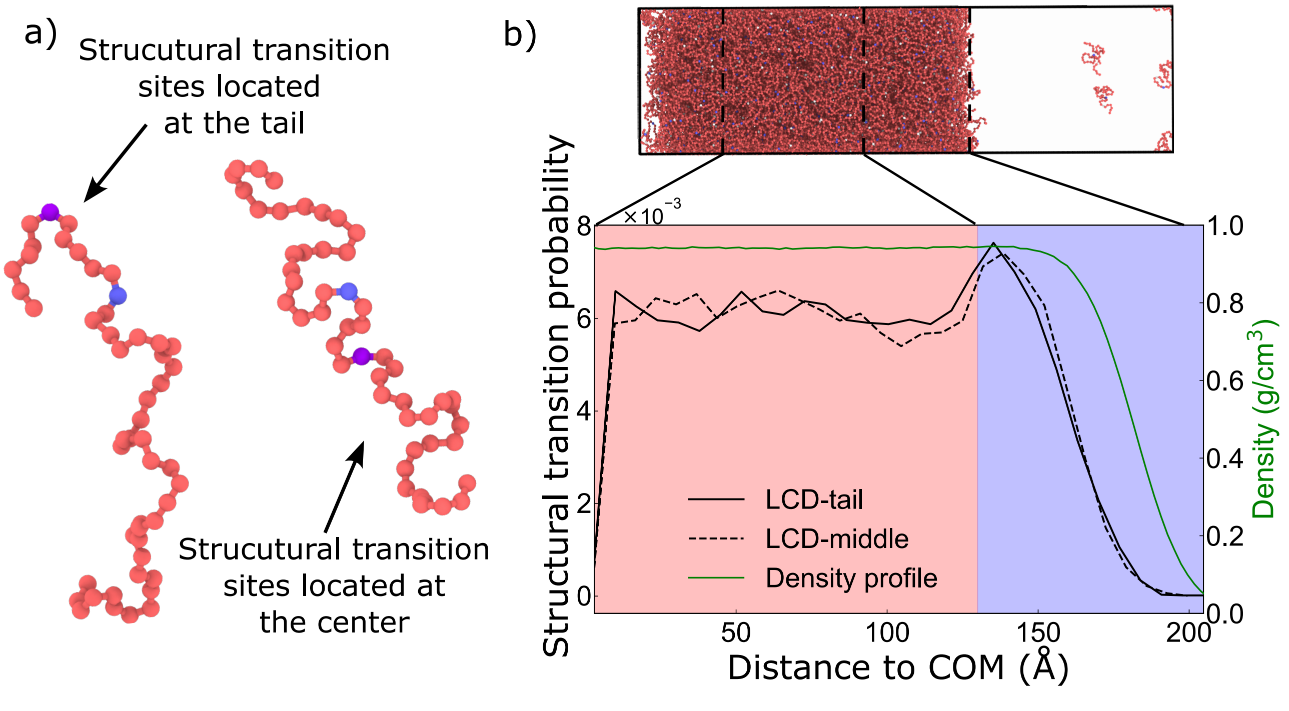


Figure S8: (a) Schematic representation of two LCDs. On the left side the structural transition sites are located in the tail (LCD-tail), while on the right they are located at the center (LCD-center). (b) Structural transition probability of LCD-tail (solid black line) and LCD-center (dashed black line) with the density profile of a condensate (green) as a function of the distance to the center of mass (COM) of the biggest cluster. At the top of the graph, a render of the slab condensate is shown, with the interface and bulk limits indicated by dashed lines.

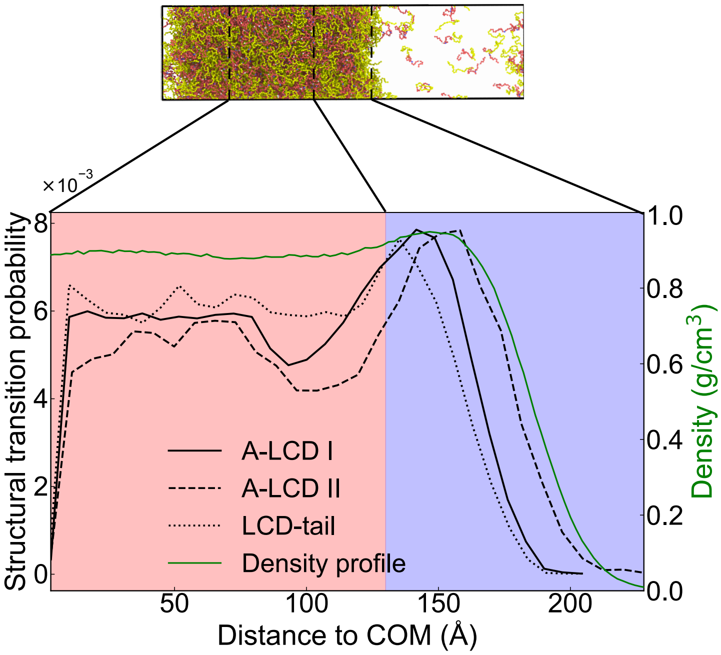


Figure S9: (a) Schematic representation of two LCDs. On the left side the structural transition sites are located in the tail (LCD-tail), while on the right they are located at the center (LCD-center). (b) Structural transition probability of LCD-tail (solid black line) and LCD-center (dashed black line) with the density profile of a condensate (green) as a function of the distance to the center of mass (COM) of the biggest cluster. At the top of the graph, a render of the slab condensate is shown, with the interface and bulk limits indicated by dashed lines.

The first 10% of the trajectory was discarded to eliminate initial equilibration effects. All statistical analysis was performed on the remaining production trajectory.

Pressure tensor components in molecular dynamics simulations exhibit temporal correlations. To properly estimate uncertainties, a block averaging procedure was employed.

Let the production trajectory contain $N$ samples. The data were divided into $M$ contiguous blocks of equal size:

$$n_b = \left\lfloor \frac{N}{M} \right\rfloor, \tag{S13}$$

where $n_b$ is the number of samples per block. For each block $i$, the block-averaged surface tension is:

$$\gamma_i = \frac{1}{n_b} \sum_{t \in \text{block } i} \gamma(t). \tag{S14}$$

The overall mean surface tension is:

$$\bar{\gamma} = \frac{1}{M} \sum_{i=1}^{M} \gamma_i. \tag{S15}$$

The variance of the block means is:

$$\sigma^2_{\text{block}} = \frac{1}{M-1} \sum_{i=1}^{M} (\gamma_i - \bar{\gamma})^2, \tag{S16}$$

and the standard error of the mean surface tension is:

$$\sigma_{\bar{\gamma}} = \sqrt{\frac{\sigma^2_{\text{block}}}{M}}. \tag{S17}$$

This approach accounts for time correlation provided the block length exceeds the autocorrelation time of the observable [8].

#### SXIII. INCREASED CONTRAST BETWEEN THE HYDROPHOBIC AND HYDROPHILIC

As commented in the main text, increasing the contrast between the hydrophobic and hydrophilic halves of the molecule further enhances the probability of structural transitions at the condensate interface. These results can be seen in Fig. S9. As the interaction disparity grows, hydrophobic residues preferentially orient toward the interface, promoting local ordering and β-sheet formation. Consequently, the location of the maximum transition probability shifts closer to the interface compared with the bulk. These results indicate that amphiphilic asymmetry not only increases interface-specific transitions but also drives their spatial localization toward the interface.


[1] J. S. Rowlinson and B. Widom, *Molecular theory of capillarity*. Courier Corporation, 2013.
